# Colour Constancy: Colour Interaction between Local Surround and Illumination in Virtual Reality Scenarios

**DOI:** 10.64898/2026.02.12.705534

**Authors:** Raquel Gil Rodríguez, Laysa Hedjar, Betül Kilic, Karl R. Gegenfurtner

## Abstract

In our study, we used virtual reality to investigate how the colour of an object’s surroundings influences colour constancy. Using Unreal Engine, we manipulated lighting and object properties in computer-generated scenes illuminated by five different light sources and presented them through an HTC Vive Pro Eye virtual reality headset. Participants assessed colour constancy by selecting the object that best matched a neutral reference from among five differently coloured options within the scene. Our results demonstrated a significant decline in colour constancy performance when the illuminant colour was in the opposite direction to that of the local surround, highlighting the interactive effects of surround colour and illumination.

## 1. Introduction

Colour shapes the way we experience the world. It helps us identify objects, navigate our surroundings, and even form lasting memories Gegenfurtner and Rieger (2000); Wichmann et al. (2002). Yet the colours we perceive are astonishingly stable, even under dramatically changing illumination. A sheet of paper, for example, appears white whether it is lit by a warm desk lamp or the fading light of sunset. This remarkable ability, known as colour constancy, allows the visual system to infer the true surface reflectance of objects across diverse lighting conditions Helmholtz (1867, 1910); Hering (1920); Judd (1940); von Kries (1923). Early investigations debated the mechanisms driving this stability, but researchers consistently found that constancy is strongest in well-structured scenes with a wide field of view Katz (1911).

A key breakthrough in quantifying colour constancy came with the introduction of the colour constancy index (CCI) Arend and Reeves (1986), which ranges from 0 (no constancy) to 1 (perfect constancy). Initial studies often reported surprisingly low CCIs, sometimes only 20%, yet methodological advances have steadily pushed reported constancy levels higher, frequently exceeding 80% under controlled conditions Foster (2011); Witzel and Gegenfurtner (2018). This improvement is particularly evident when scenes are illuminated naturally and dominated by a single, well-defined light source Gegenfurtner et al. (2024); Hansen et al. (2007); Hurlbert et al. (2019); Kraft and Brainard (1999); Olkkonen et al. (2010); Radonjić et al. (2015, 2018).

More recent evidence suggests that colour constancy is further enhanced when experimental tasks closely resemble real-world viewing conditions Gegenfurtner et al. (2024). Traditionally, achieving this level of realism required carefully controlled physical setups, which, while providing ecological validity, often limited experimental flexibility. The advent of virtual reality (VR) and modern 3D rendering technologies now offers a solution: researchers can create immersive, naturalistic environments while maintaining precise experimental control Cipresso et al. (2015); Gil Rodriguez et al. (2024); Hou et al. (2025a,b); Shapiro and LoPrete (2020); Wiesing et al. (2020); Yaremych and Persky (2019). This combination of realism and control opens exciting new avenues for investigating human colour perception under conditions that closely mirror everyday life. This is what we did in our previous experiments, where we combined the controlled colour characterisation of new head-mounted displays (HMDs) with the photorealistic rendering of current gaming engines like Unreal Engine Gil Rodriguez et al. (2022, 2024).

In our previous study, we used virtual reality to evaluate colour constancy in both indoor and outdoor scenes, achieving high levels of constancy similar to those observed in real-world immersive experiments Gegenfurtner et al. (2024); Gupta et al. (2020); Kraft and Brainard (1999); Mizokami et al. (2004); Morimoto et al. (2017); Olkkonen et al. (2010); Pearce et al. (2014). Our results highlight the advantages of VR for these studies, including precise cue control, accurate colour reproduction, and preserved immersion. We investigated key mechanisms underlying colour constancy:

- Local surround Wallach (1948): An object’s immediate surroundings strongly influence colour constancy, with effects amplified when the surround contrasts with the illuminant, consistent with previous findings on the role of local contrast in multi-surface scenes Brown and MacLeod (1997); Hurlbert and Wolf (2004); Morimoto et al. (2024a, 2021, 2017); Olkkonen and Ekroll (2016); Olkkonen et al. (2010); Toscani et al. (2025).
- Maximum fluxLand (1986); Land and McCann (1971): The brightest object in a scene did not significantly affect colour constancy in our experiments, although trends suggest it may contribute under certain conditions, and its effectiveness may depend on object prominence, scene structure, and display dynamic range Morimoto et al. (2026).
- Spatial mean Brainard and Wandell (1986); Buchsbaum (1980): The average colour of a scene remains an important factor. Our findings indicate that observers rely on some form of scene segmentation rather than a simple pixel-wise computation, and that long-range chromatic context and object reflectances influence constancy, especially when the spatial mean cue is manipulated.

One of the most striking findings involved the local surround mechanism. Colour constancy decreased significantly when this cue was silenced, with the effect being particularly pronounced under the green illuminant. We hypothesise that this arises from an interaction between the illuminant and the immediate surround, which in our experiment was rose-coloured, enhancing contrast effects on the perceived target colour. Building on these previous findings Gil Rodriguez et al. (2024), the present study explores how local surround cues interact with illuminant colour to shape human colour constancy in immersive virtual environments. In our experiments, participants viewed objects in both indoor and outdoor scenes under five different illuminants and four surround colours. We compared performance between Baseline conditions, where all cues were accessible, and Local Surround Silencing conditions, where the constancy cue from the immediate chromatic context was removed. Our analyses revealed that colour constancy is highly context-dependent: indoors, constancy remained robust under Baseline conditions, while outdoor scenes demonstrated greater vulnerability to local and global cue interactions. We found that keeping local surroundings constant across illuminations consistently diminished constancy, with larger deficits evident for illuminants opposing the surround color and in more complex outdoor settings. These findings illustrate how the visual system adeptly integrates local and global information, utilising chromatic context to stabilise perceived object colour.

## 2. Methodology

### 2.1. Observers

Two authors and 13 naïve observers participated in the study, with 8 completing the experiment in an indoor scene and another 8 in an outdoor scene (one participant completed the experiment in both scenes). All participants had normal colour vision, which was confirmed using the Ishihara Colour Test. This study was approved by the ethical commission of the Department of Psychology at Justus-Liebig-Universität Giessen (LEK 2020-0015). Excluding one author, the observers were students recruited from Justus-Liebig-Universität Giessen, and all were fluent English speakers. They all completed the experiment in several sessions.

### 2.2. Equipment

The experiments were conducted on a desktop computer running Windows 10, equipped with an Intel® Core™ i9-9900K processor (3.60GHz), 128GB of RAM, and an NVIDIA GeForce RTX 4090 graphics card. Visual stimuli were presented using an HTC Vive Pro Eye VR headset, with the virtual environments developed and rendered in Unreal Engine v4.27. Data analysis was performed using MATLAB R2024b Inc. (2024) and *RStudio* R Core Team (2025); Team (2021).

### 2.3. Colour Characterisation

Accurate colour characterisation of the display is essential in colour perception research Brainard et al. (2002). We followed the same characterisation procedure described in Díaz-Barrancas et al. (2024). For this, we measured 52 values per primary (red, green, and blue), along with a grey ramp consisting of 52 additional measurements. All RGB values were rendered in Unreal Engine using the same engine settings as in the experiment. The resulting model comprises a linear transformation, represented by a matrix whose columns correspond to the XYZ measured values of the maximum intensities in each primary, and a non-linearity based on the grey ramp, implemented using three 1D look-up tables (LUTs). All measurements were obtained using a Konica Minolta CS-2000A spectroradiometer Konica Minolta (2022). The model was validated using 100 random RGB values and their corresponding measured XYZ values, resulting in an average error, Δ*E*00 Sharma et al. (2005), of less than 1, indicating perceptual equivalence.

### 2.4. Stimulus

#### 2.4.1. Two rendered scenarios

Two scenes were rendered for the experiment (Figure 1). The first was an indoor, office-like environment featuring furniture such as a chair, an armchair, a table, and shelves. The room was lit by a single *point light* positioned above the observer, in the centre of the ceiling. This light emits evenly in all directions but is not visible to the observer.

**Fig. 1.**
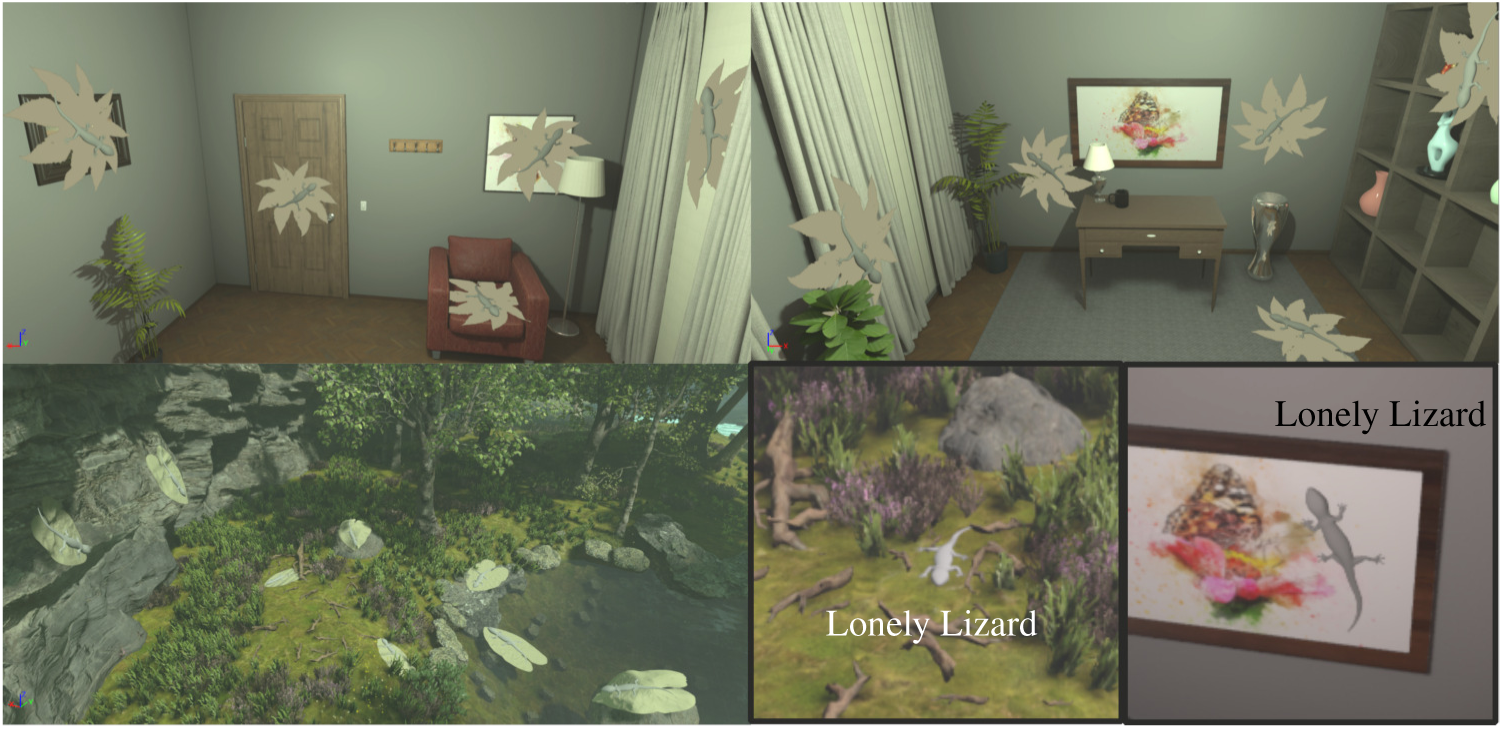
Two 3D-rendered scenes are presented under green illumination. The top row displays two images of the indoor scene—one from the back view and one from the front. In both images, leaves with lizards on them indicate the 10 locations used in the selection task. The bottom row shows the outdoor scene, featuring 8 different lizards positioned at the various locations considered for the experiment. The bottom-right image displays the ‘Lonely Lizard’ used in both the indoor and outdoor scenes under neutral illuminant.

The second scene was an outdoor, forest-like environment with elements such as trees, cliffs, grass, flowers, and a lake. It was illuminated using a *directional light*, simulating sunlight, which casts light consistently from one direction. In addition, a *skylight* component was used to softly illuminate the scene using light and colour captured from the sky, helping to reduce harsh shadows and enhance reflections. Both the directional light and the skylight shared the same colour for consistency.

#### 2.4.2. Illuminant and local surround colours

Five distinct illuminants were tested, each carefully selected to explore a range of chromatic shifts. The (*x*, *y*) chromaticity coordinates for these illuminants were based on those used in Aston et al. (2019); Gil Rodriguez et al. (2024), ensuring consistency with previous research. The tested illuminants and their (x,y) coordinates included: neutral (0.31, 0.33), blue (0.25, 0.26), yellow (0.39, 0.39), green (0.30, 0.38) and red (0.32, 0.26). To convert the chromaticity values to RGB input values for the engine, we set the luminance to *Y* = 30 cd/m^2^ and use the previously defined colour characterisation for this conversion Díaz-Barrancas et al. (2024).

In addition to the illuminant colours, we manipulated the colour of the immediate local surround to assess its impact on colour constancy performance. These surround colours were defined within the CIELAB colour space and positioned diagonally to the illuminant colours. We named them *Khaki, Rose, Purple,* and *Teal*, as shown in Figure 2.

**Fig. 2.**
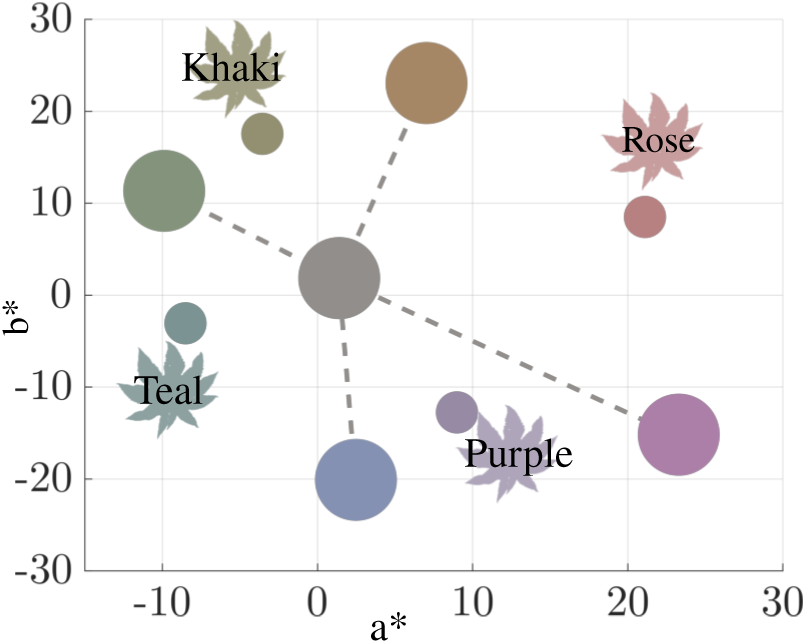
Chromaticity values (a*, b*) in CIELAB colour space of the five illuminants (big circles) and the four local surround colours (small circles) with the corresponding surround shape (leaf). The colours used in this plot are for visualisation purposes only.

#### 2.4.3. Reference object and competitors

The reference object is an achromatic lizard, with its surface reflectance set to match the chromaticity of the VR headset’s white point. This reference, referred to as the ‘Lonely Lizard’, is rendered in Unreal Engine using a simple material setup: the base colour corresponds to the calibrated achromatic value, while the metallic and specular parameters are set to 0, and the roughness is set to 1, creating a matte-like surface appearance (Figure 1, bottom right).

Under each illuminant, we defined five coloured lizards for our selection task. The method used to determine these colours follows the same procedure described in Radonjić et al. (2015); Radonjić et al. (2015). Our *reflectance* value corresponds to the colour of the reference object, the *Lonely Lizard*. Next, we define the *tristimulus* value, which is computed such that its reflected light under the specific coloured illumination matches the reflected light of the *reflectance* value under the neutral illumination. Once these two anchor points are defined, we generate two intermediate samples positioned between them in CIELAB space. Additionally, we compute an *overconstant* value, which lies along the same axis as the line from the *reflectance* to the *tristimulus*, but in the opposite direction and at 25% of the total distance between the two anchor points.

Note that under the neutral illuminant, the *reflectance* and *tristimulus* values are identical. In this case, we defined two samples taken from the competitor list under the blue illuminant and two under the yellow illuminant. From each illuminant, one of these samples was the computed *tristimulus* value. The second was positioned between this *tristimulus* value and the *reflectance* value, which is the same across all illuminants.

#### 2.4.4. Baseline and Local Surround suppression conditions

We set up two conditions: 1) *Baseline*, and 2) *Surround Silencing*. In the Baseline condition, which serves as the control in which all constancy cues are available, the lizards are placed on top of coloured leaves (see Section 2.4.2). The leaves are treated as matte surfaces that interact with the illumination and visual stimulus differently depending on the colour of the illuminant, as shown in the top row in Figure 3. In contrast, in the Surround Silencing condition, we treat the leaves as self-emissive surfaces, meaning the visual stimulus remains constant and does not change with illumination, as shown in the bottom row in Figure 3.

**Fig. 3.**
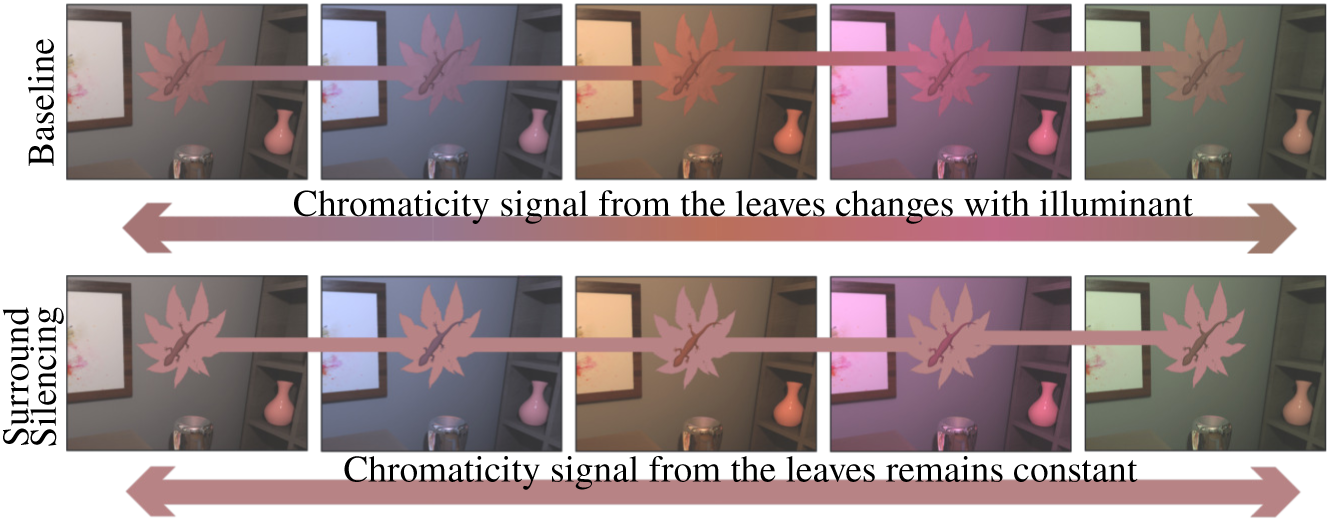
At the top: The indoor environment under five different types of illumination (neutral, blue, yellow, red, and green). At the bottom: The same environment illustrates how local surround suppression was implemented. The chromaticity of the reflected light from a leaf beneath the target object (a lizard) remains constant despite changes in illumination.

### 2.5. Experimental Procedure

#### 2.5.1. Data Collection

Participants were required to complete an object-selection task under varying illumination conditions. The objective was to identify a specific reference object, referred to as the ‘Lonely Lizard’, under different illuminations. To familiarise participants with the reference object, the Lonely Lizard was first presented under a neutral illuminant, allowing observers to observe its baseline appearance without the influence of colour shifts in illumination (Figure 1, bottom right).

After participants viewed the Lonely Lizard under neutral illumination, the illuminant in the scene changed to one of the coloured illuminants or remained the same, and the Lonely Lizard disappeared. Over the following 90 seconds, participants were instructed to explore the scene in search of a small ‘gnome’ character that floated around randomly and occasionally disappeared for brief intervals. This task encouraged active visual engagement with the environment and promoted adaptation to the new lighting. After the 90 seconds, the gnome disappeared, and five lizards appeared around the scene; see Figure 1 for an overview of the possible lizard positions. Participants’ task was to locate and identify the same Lonely Lizard under the new illumination using the controller. The Lonely Lizard itself was achromatic; however, this was not explicitly stated to participants to avoid biasing their selection process. Instead, they were instructed to search for the same Lonely Lizard shown previously, now illuminated in the scene by a different colour. This was repeated for a total of 30 trials per illuminant (see Figure 4). The task was always blocked by illumination and leaf colour, aligning with the procedure in our previous work Gil Rodriguez et al. (2024).

**Fig. 4.**
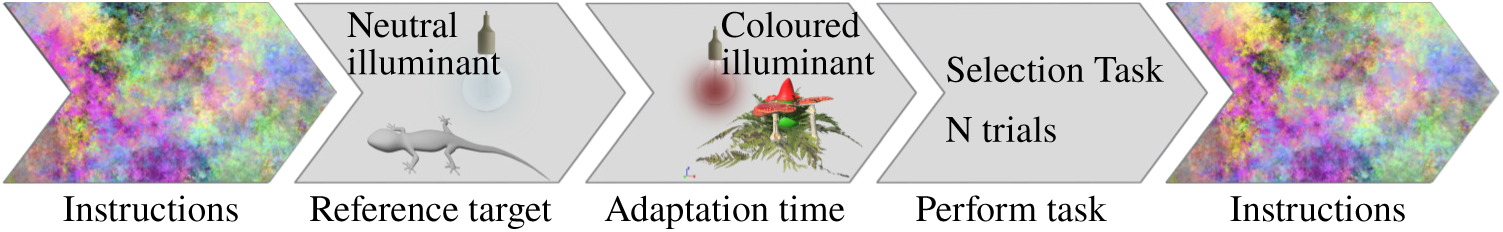
Colour constancy pipeline: 1) participants view instructions in a room with pink noise textures on the walls, 2) they are shown a reference lizard (the ‘Lonely Lizard’) under a neutral illuminant, 3) after adapting for 1.5 min under a coloured illuminant, 4) participants are presented with four competitors and the reference target and asked to select the one matching the reference (this is repeated across several trials), and 5) participants return to the pink noise textured room.

Participants completed the Baseline conditions with four different local surround (i.e., leaf) colours, as well as the Local Surround Suppression condition under the same four leaf colours, under all five illuminants. This resulted in a total of 40 different experimental conditions per scene. Eight observers completed these conditions for the indoor scene and 8 for the outdoor scene. They performed the tasks across multiple sessions, also ensuring that they did not wear the VR headset for more than 30 minutes at a time.

Participants conducted the experiments while seated in a chair positioned at the centre of the room. Although they were free to move, the task did not require it. Their movements were mostly limited to rotating the chair.

#### 2.5.2. From Selection Task to Individual Perceptual Space

We utilise the method described in Radonjić et al. (2015); Radonjić et al. (2015), which is an adaptation of the original maximum likelihood difference scaling (MLDS) method described in Maloney and Yang (2003), to infer participants’ internal perceptual representations based on their choices in the selection task experiment.

The method described in Radonjić et al. (2015) employs a paired comparison approach, while our task is a multiple-choice selection. This requires us to transform the observer’s selections into paired comparisons. We accomplish this by defining all ten pairwise combinations of the five competitors. During each trial, participants are shown all five competitors simultaneously, and we record a vote for the chosen competitor in each pair where it appears. This means that some of the pairs might have no votes.

Once the data is reformatted to reflect paired comparison responses, the algorithm estimates the positions of the five competitors within a one-dimensional perceptual space, along with the participant’s *inferred match*. The relative order of the competitors is preserved throughout the estimation, with the first competitor always fixed at position 0. Each competitor’s location is assumed to follow a Gaussian distribution with a standard deviation of *σ* = 0.1, and a minimum distance from each other of 0.025 units.

The final step involves defining the inferred match from the one-dimensional perceptual space to the CIELAB colour space. We achieve this by maintaining the same ratios to the CIELAB colour space, as described in Radonjić et al. (2015).

#### 2.5.3. Colour Constancy Index (CCI)

Let 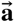 denote the vector from the participant’s match under the neutral illuminant to the ideal match under the coloured illuminant. We define 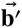 as the projection of the vector from the participant’s ideal coloured match to their selected coloured match onto the direction of 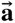. The Colour Constancy Index (CCI) is then given by

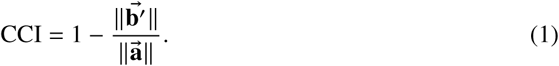

A CCI of 1 indicates perfect colour constancy, while a CCI of 0 indicates no constancy. All distances are computed within the CIELAB colour space to ensure perceptual relevance. Figure 5 illustrates an example of the CCI computation, where the participant’s CCI is approximately 0.45.

**Fig. 5.**
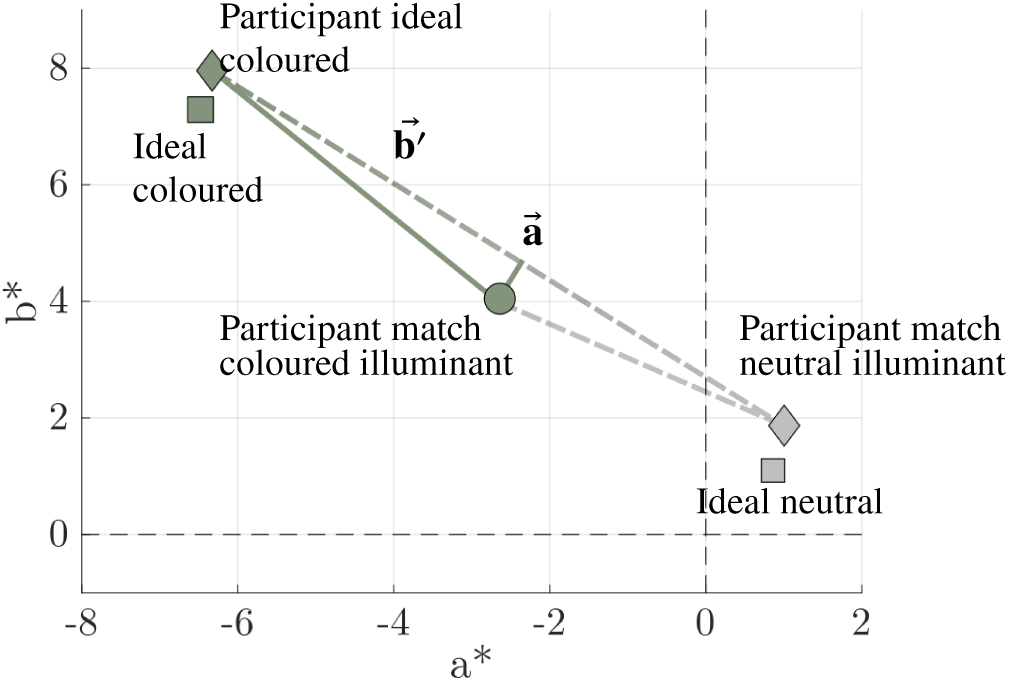
Colour constancy index definition on the chromatic axes of CIELAB colour space.

Each *Participant’s match neutral illuminant*, grey diamond in Figure 5, is defined as the average of their responses across the four Local Surround colours within each experimental condition and scene combination. Thus, for each participant, we obtained one neutral illuminant estimate for the Indoor scene and Baseline condition, as well as for the Indoor scene and Surround Silencing condition. The same applies to the Outdoor scene. The neutral illuminant condition, along with participants’ match variability, is reported in Appendix C.

We also define an error metric, the *signed error* (Δ*SE*), as follows:

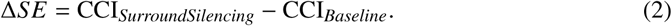

A Δ*SE* value of 0 means that the CCI in the Surround Silencing and Baseline conditions were the same. The larger the Δ*SE*, the larger the difference between the CCIs of the two conditions. A negative value means the CCI in the Suppression condition was smaller than in the Baseline, while a positive value means the reverse. We use it to analyse and plot the data.

#### 2.5.4. Data Analysis

We analyse the participants’ performance using an *ANOVA* within a *linear mixed-effects model* (LME) Bates et al. (2015). In this model, we treat participants as a random effect and the different factors and their interactions as fixed effects, as listed below, with CCI as the response variable. This approach enables us to assess the impact of each factor while controlling for random variation due to individual differences among subjects.

- **Surround colour:** Khaki, Rose, Purple, and Teal.
- **Scene:** Indoor and Outdoor.
- **Experimental condition:** Baseline and Surround Silencing.
- **Illuminant colour:** Blue, Yellow, Green, and Red.

For those fixed factors with more than two levels, and where a significant difference is found, we will perform *post hoc tests* using estimated marginal means (*emmeans*) and contrasts using pairwise comparisons within each level, and applying Holm correction.

We assessed the model’s confidence in estimating each participant’s inferred match using the Pearson coefficient. This confidence metric reflects how well the model fits an individual’s data, serving as an indicator of its reliability in capturing the structure of perceptual responses. To ensure the robustness and interpretability of our results, we excluded one participant under a specific condition whose model confidence fell below 0, representing only 0.2% of the total data. Low-confidence cases may result from inconsistent response patterns or situations in which the model failed to describe the participant’s responses accurately. Overall, 88% of the model confidence responses were above 0.95.

## 3. Results

Overall, the statistical analysis showed significant main effects of Illuminant, Surround colour, Scene, and Experimental condition. Crucially, two three-way interactions were significant: Illuminant × Surround colour × Scene and Illuminant × Surround colour × Experimental condition. Silencing the local surround significantly reduced colour constancy in both scenes, with a larger decrease in Outdoor than in Indoor. Illuminant-dependent differences were minimal under Baseline conditions but emerged clearly under Surround Silencing. These effects depended systematically on the chromatic relationship between the illuminant and the local surround: illuminants that were chromatically *Opposing* to the surround produced larger reductions in colour constancy than *Neighbouring* illuminants, especially in the Outdoor scene. An analysis of this complementary chromatic relationship showed a strong main effect of Opposing versus Neighbouring illuminants and interactions with Scene and Experimental condition.

### Main Effects and Interactions

We selected a linear mixed-effects model that included all two-way and three-way interactions of all factors *Surround colour*, *Illuminant colour*, *Experimental condition*, and *Scene* as fixed effects, and *Participant* as a random effect. We excluded the four-way interaction based on statistical evidence. A likelihood-ratio test comparing the simpler model against the full model with the four-way interaction showed no improvement in fit, *p* = 0.906. Information criteria further supported this, both AIC −358.30 and BIC −110.60 favoured the simpler model. In addition, model diagnostics confirmed that the residuals were approximately normally distributed, as indicated by the Shapiro-Wilk SHAPIRO and WILK (1965) test, *p* = 0.105.

Analysis of the linear mixed-effects model using a Type III Analysis of Variance with Satterthwaite’s approximation Satterthwaite (1946) revealed several significant main effects and interactions (Table 1). There was a significant main effect of *Illuminant* (*F*(3, 499.57) = 8.08, *p* < 0.001, 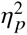 = 0.052), and a smaller but reliable effect of *Surround colour* (*F*(3, 499.88) = 2.97, *p* = 0.031, 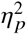 = 0.022). Both Scene (*F*(1, 50.24) = 16.00, *p* < 0.001, 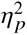 = 0.233) and *Experimental condition* (*F*(1, 499.92) = 62.54, *p* < 0.001, 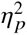 = 0.124) showed robust main effects, indicating substantial differences in colour constancy across scene types and between Baseline and Surround Silencing conditions. See Figures 12 and 13 for the Indoor and Outdoor scenes, respectively. These figures show CCI values for each local surround colour under the four coloured illuminants. For each illuminant, the paired bar plots compare the Baseline and Surround Silencing conditions.

**Table 1.** Type-III ANOVA results (Satterthwaite approximation) for the linear mixed-effects model.

| | df | F | p-value | $\eta_p^2$ |
| --- | --- | --- | --- | --- |
| Illuminant | 3 | 8.08 | $2.91 \times 10^{-5}$ *** | 0.052 |
| Surround colour | 3 | 2.97 | 0.0313 * | 0.022 |
| Scene | 1 | 16.00 | $2.08 \times 10^{-4}$ *** | 0.233 |
| Experimental condition | 1 | 62.54 | $1.69 \times 10^{-14}$ *** | 0.124 |
| Illuminant × Surround colour | 9 | 5.88 | $8.04 \times 10^{-8}$ *** | 0.109 |
| Illuminant × Scene | 3 | 2.14 | 0.0941 | 0.009 |
| Surround colour × Scene | 3 | 0.76 | 0.515 | 0.006 |
| Illuminant × Experimental condition | 3 | 4.25 | 0.00558 ** | 0.029 |
| Surround colour × Experimental condition | 3 | 1.36 | 0.256 | 0.010 |
| Scene × Experimental condition | 1 | 5.33 | 0.0213 * | 0.010 |
| Illuminant × Surround colour × Scene | 9 | 2.46 | 0.00956 ** | 0.044 |
| Illuminant × Surround colour × Experimental condition | 9 | 2.11 | 0.0274 * | 0.039 |
| Illuminant × Scene × Experimental condition | 3 | 0.31 | 0.819 | 0.005 |
| Surround colour × Scene × Experimental condition | 3 | 0.24 | 0.868 | 0.002 |

Several interaction terms were also significant. In particular, *Illuminant* interacted strongly with *Surround colour* (*F*(9, 499.56) = 5.88, *p* < 0.001, 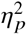 = 0.109) demonstrating that the effect of the illuminant depended on the chromatic properties of the local surround. A significant *Illuminant* × *Experimental condition* interaction was also observed (*F*(3, 499.59) = 4.25, *p* = 0.0056, 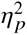 = 0.029), indicating that illuminant effects differed between the Baseline and Surround Silencing conditions. In addition, *Scene* interacted with *Experimental condition* (*F*(1, 499.91) = 5.33, *p* = 0.021, 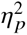 = 0.010).

Two three-way interactions reached significance. The interaction between *Illuminant*, *Surround colour*, and *Scene* was reliable (*F*(9, 499.56) = 2.46, *p* = 0.010, 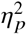 = 0.044), as was the interaction between *Illuminant*, *Surround colour*, and *Experimental condition* (*F*(9, 499.55) = 2.11, *p* = 0.027, 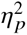 = 0.039). All remaining interaction terms were non-significant (all *p* > 0.24, 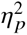 ≤ 0.010).

### Pairwise Contrasts

To quantify how colour constancy changed when the local surround cue was silenced, we estimated the simple effect of *Experimental condition* separately for each *Scene* using estimated marginal means. Colour constancy decreased significantly under Surround Silencing in both environments. In Indoor scenes, CCI decreased by 0.085 (*t*(440) = 4.11, *p* < 0.0001), whereas in Outdoor scenes the decrease was substantially larger at 0.147 (*t*(441) = 7.02, *p* < 0.0001). These results confirm that silencing local information has a stronger detrimental effect on colour constancy in Outdoor than in Indoor scenes; see Figure 6. The remaining two-way interactions are reported in the Appendix B.1.

**Fig. 6.**
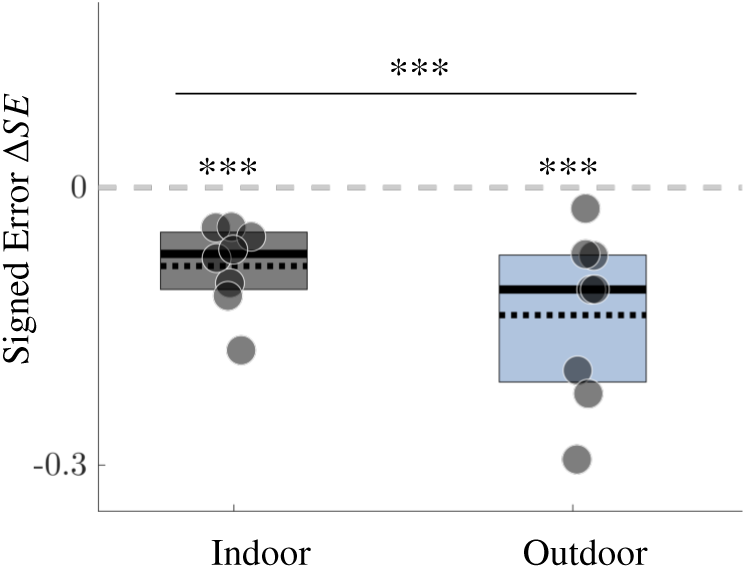
Signed error Δ*SE* (Silencing - Baseline) for each participant (dot), comparing the two conditions in both scenes. Negative values indicate reduced performance when the local surround cue is suppressed.

To unpack the significant 3-way interaction between *Illuminant*, *Surround colour*, and *Scene*, we examined pairwise illuminant contrasts separately for each Surround colour within Indoor and Outdoor scenes using estimated marginal means (Holm-corrected). In Indoor scenes, illuminant effects were generally weak and selective. No significant illuminant differences were observed for the Khaki or Teal surrounds (all adjusted *p* = 1.00). In contrast, for the Purple surround, Blue elicited higher CCI values than Green (0.215, *t*(440) = 3.68, *p* = 0.0016), with a similar but non-significant trend relative to Yellow (0.147, *p* = 0.060). No reliable illuminant differences were found for the Rose surround in Indoor scenes. In Outdoor scenes, illuminant effects became more pronounced and depended strongly on Surround colour. For the Khaki surround, Green yielded higher CCIs than both Red (0.158, *t*(440) = 2.71, *p* = 0.035) and Yellow (0.167, *t*(440) = 2.87, *p* = 0.026). In the Purple surround, Blue exceeded both Green (0.185, *t*(440) = 3.05, *p* = 0.012) and Yellow (0.240, *t*(440) = 4.11, *p* = 0.0003), and Red was also higher than Yellow (0.174, *t*(440) = 2.99, *p* = 0.012). For the Rose surround in Outdoor scenes, the pattern reversed: Blue produced lower CCI than Red (−0.190, *t*(440) = −3.26, *p* = 0.006), and Green was also lower than both Red (−0.220, *t*(440) = −3.70, *p* = 0.0014) and Yellow (−0.169, *t*(440) = −2.84, *p* = 0.0187). Finally, for the Teal surround, Blue exceeded Red (0.192, *t*(440) = 3.29, *p* = 0.0047) and Yellow (0.225, *t*(440) = 3.79, *p* = 0.0010), and Green was higher than both Red (0.165, *t*(440) = 2.82, *p* = 0.015) and Yellow (0.198, *t*(440) = 3.33, *p* = 0.0047). Together, these results show that illuminant-dependent differences in colour constancy were minimal in Indoor scenes but emerged robustly in Outdoor scenes, with the direction and magnitude of the effect strongly modulated by Surround colour.

A second significant 3-way interaction was observed between *Illuminant*, *Surround colour*, and *Experimental condition* (*F*(9, 499.55) = 2.11, *p* = 0.027, 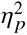 = 0.039). Pairwise illuminant contrasts were therefore examined separately for each Surround colour under Baseline and Surround Silencing conditions. Under the Baseline condition, illuminant differences were uniformly small, and none of the contrasts reached significance for any Surround colour (all adjusted *p* ≥ 0.38), indicating little sensitivity to illuminant identity when the local surround cue was available. In contrast, clear illuminant effects emerged under Surround Silencing. For the Khaki surround, Green produced a higher CCI than Red (0.177, *t*(440) = 3.03, *p* = 0.0156). In the Purple surround, a strong illuminant effects were observed: Blue exceeded Green (0.299, *t*(440) = 4.92, *p* < 0.0001) and Yellow (0.301, *t*(440) = 5.16, *p* < 0.0001), and Red also exceeded Yellow (0.182, *t*(440) = 3.12, *p* = 0.0076). For the Rose surround, Green yielded a lower CCI than Red (−0.241, *t*(440) = −4.06, *p* = 0.0003), whereas other contrasts were not significant. Finally, in the Teal surround, Blue was higher than both Red (0.186, *t*(440) = 3.18, *p* = 0.0078) and Yellow (0.270, *t*(440) = 4.62, *p* < 0.0001), and Green also exceeded Yellow (0.153, *t*(440) = 2.62, *p* = 0.036).

### Chromatic Shift and Signed Error (Δ*SE*)

We investigated whether there is a relationship between the shift in chromaticity signal of the leaf between the Baseline condition and the Surround Silencing condition (see Figure 10), and the participants’ Δ*SE* values. For each combination of Surround colour, Illuminant, and Scene, we averaged the Δ*SE* and chromaticity shift values. We then computed the Pearson correlation between these two sets of values. The analyses revealed negative correlations (Indoor: r = −0.68; Outdoor: r = −0.49), indicating that a smaller chromaticity shift corresponds to a signed error Δ*SE* closer to zero. See Figure 7.

**Fig. 7.**
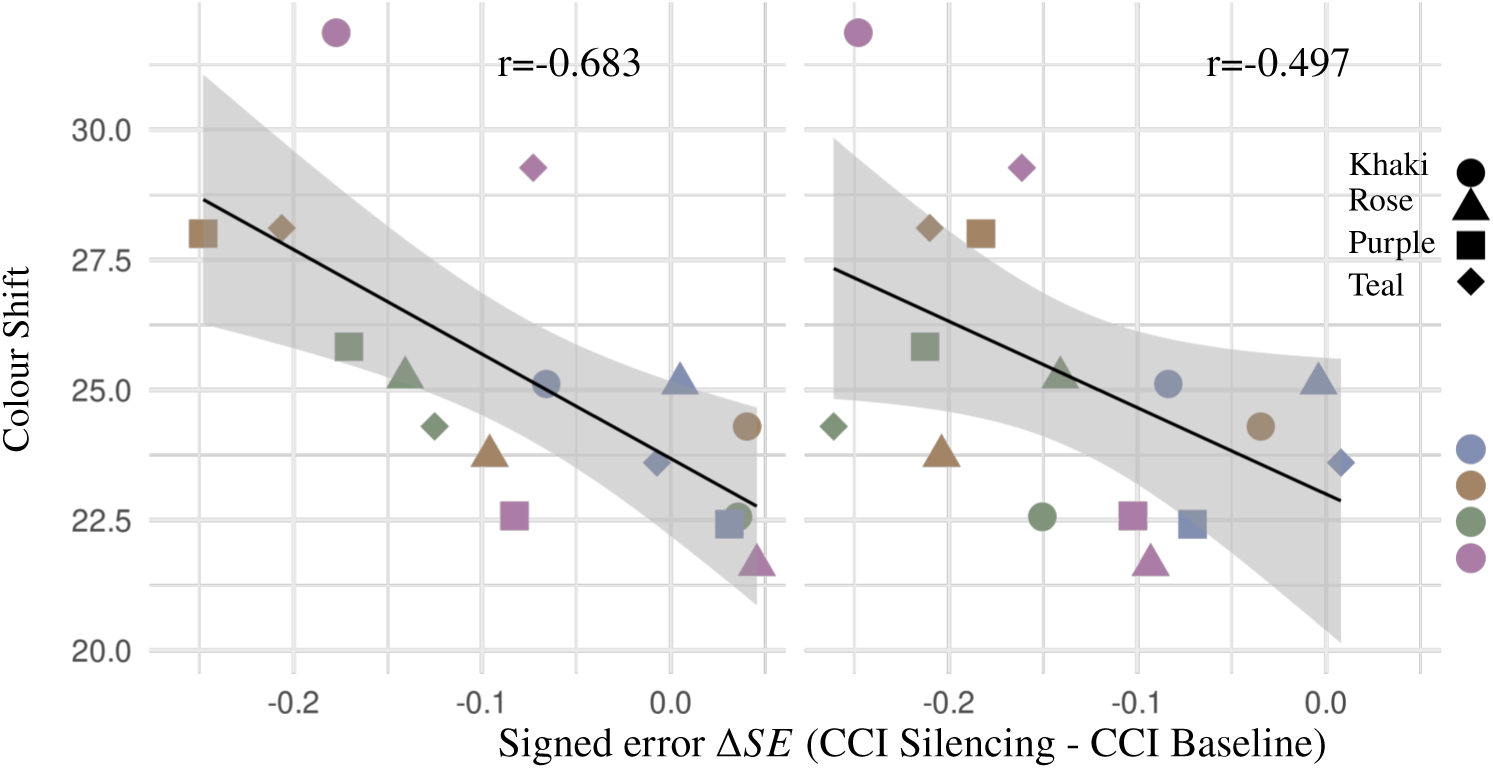
Colour shift versus signed error Δ*SE* in the Indoor and Outdoor scenes for each Surround colour (circle-khaki, triangle-rose, square-purple, and diamond-teal), shown separately for all illuminants (blue, yellow, green and red).

Figure 8 shows that the signed error, Δ*SE*, is strongly dependent on both the illuminant and the surround colour. Distinct patterns are observed when comparing indoor and outdoor scenes; specifically, in the latter, there is a more pronounced effect under the green illuminant, regardless of the surround colour used. Overall, it is clear that for each surround colour, the illuminant that produces the smallest Δ*SE* value is the one that is opposite in hue to that colour. We can visualise this interaction pattern in Figure 8. Certain combinations of surround colour and chromatic direction lead to significant performance reductions, while others result in smaller or even reversed effects. For example, under a Khaki surround with a Red illuminant, we observe the largest drop in performance, whereas with a Yellow illuminant, the effect is reversed. This pattern tends to hold for all surround colours and scenes.

**Fig. 8.**
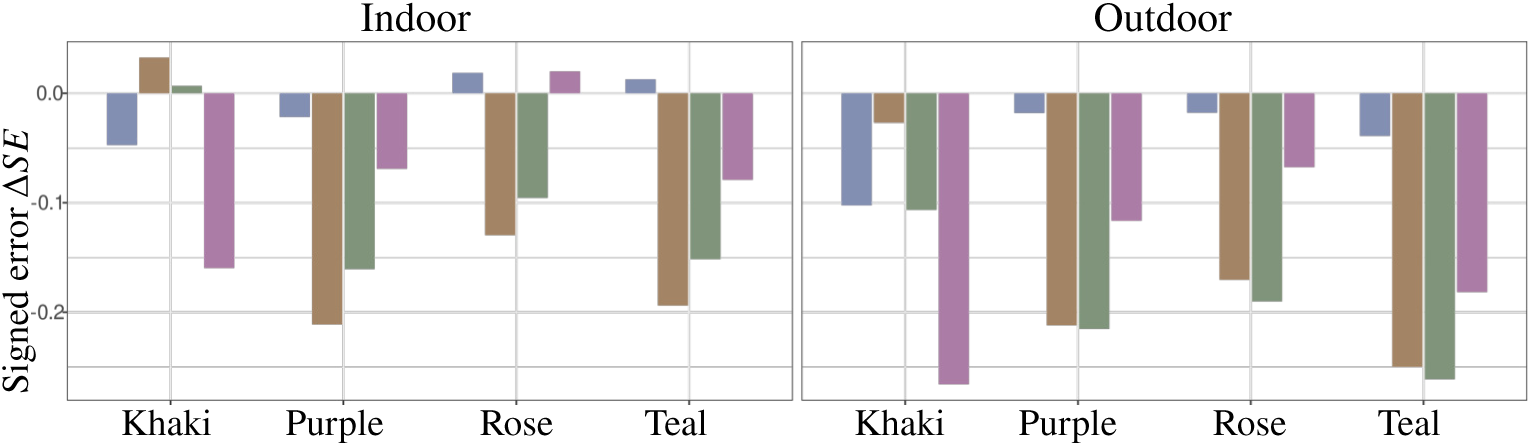
Signed error Δ*SE* in the Indoor and Outdoor scenes for each Surround colour, shown separately for all illuminants (blue, yellow, green and red) within each surround.

To characterise how the colour of the local surroundings interacts with different illumination colours, we introduce the concept of *Chromatic direction*. In our stimulus configuration, both the colours of the illuminant and the local surroundings vary along well-defined chromatic axes in the CIELAB colour space. For instance, in the case of the Khaki surround colour, we utilised Yellow and Green illuminants to define this leaf colour. We will refer to these illuminants as *Neighbouring* illuminants, while the other two, Red and Blue, will be considered as *Opposing* illuminants, see Figure 10. In the next section, we will introduce this new factor and study and analyse the colour interactions.

### 3.1. Chromatic direction: the interaction between illuminant and local surround colour

Across scenes and surround colours, colour constancy was significantly lower for Opposing than for Neighbouring illuminants, and this effect persisted across Experimental conditions. The statistical analysis reported below confirms this pattern and quantifies its interaction with Scene and Surround colour.

#### Main Effects and Interactions

A linear mixed-effects model was fitted to examine the effects of *Direction* (Neighbouring vs. Opposing), *Surround colour*, *Scene* (Indoor vs. Outdoor), and *Experimental condition* (Baseline vs. Silencing) on CCI. The model included all main effects, two-way, and three-way interactions, with a random intercept for Participant. Type III ANOVA with Satterthwaite’s approximation revealed significant main effects of *Direction*, *F*(1, 465.86) = 45.78, *p* < 0.001, 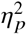 = 0.089, *Surround colour*, *F*(3, 466.20) = 3.15, *p* = .025, 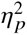 = 0.020, *Scene*, *F*(1, 44.50) = 13.88, *p* = 0.001, 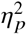 = 0.238, and *Experimental condition*, *F*(1, 466.25) = 59.36, *p* < 0.001, 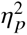 = 0.113.

Several two-way interactions were also significant. In particular, *Direction* interacted with *Surround colour*, *F*(3, 465.85) = 4.56, *p* = 0.004, 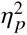 = 0.029, Scene, *F*(1, 465.86) = 10.09, *p* = 0.002, 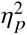 = 0.021, and *Experimental condition*, *F*(1, 465.88) = 8.39, *p* = 0.004, 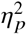 = 0.018. In addition, a significant interaction was observed between *Scene* and *Experimental condition*, *F*(1, 466.25) = 4.16, *p* = 0.042, 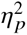 = 0.009. All remaining two-way interactions were not significant (*p* > 0.24, 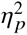 ≤ 0.009), and none of the three-way interactions reached statistical significance (all *p* > 0.15, 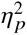 ≤ 0.007).

#### Pairwise contrasts

Because *Direction* participated in multiple significant interactions, its main effect was not interpreted in isolation. Estimated marginal means were therefore computed, and Neighbouring-Opposing contrasts were performed using Holm correction. Averaged across Scene and Experimental condition, Direction effects were significant for Purple (0.163, *t*(466) = 5.42, *p* < 0.0001), Rose (0.096, *t*(466) = 3.20, *p* = 0.0015), and Teal surrounds (0.132, *t*(466) = 4.40, *p* < 0.0001), but not for Khaki (0.015, *t*(466) = 0.50, *p* = 0.62).

**Table 2.** Type-III ANOVA results (Satterthwaite approximation) for the linear mixed-effects model.

| | F | p-value | $\eta_p^2$ |
| --- | --- | --- | --- |
| Direction | 45.78 | < 0.001 | 0.089 |
| Surround colour | 3.15 | 0.025 | 0.020 |
| Scene | 13.88 | 0.001 | 0.238 |
| Experimental condition | 59.36 | < 0.001 | 0.113 |
| Direction × Surround colour | 4.56 | 0.004 | 0.029 |
| Direction × Scene | 10.09 | 0.002 | 0.021 |
| Surround colour × Scene | 0.83 | 0.479 | 0.005 |
| Direction × Experimental condition | 8.39 | 0.004 | 0.018 |
| Surround colour × Experimental condition | 1.38 | 0.247 | 0.009 |
| Scene × Experimental condition | 4.16 | 0.042 | 0.009 |
| Direction × Surround colour × Scene | 1.02 | 0.383 | 0.007 |
| Direction × Surround colour × Experimental condition | 1.38 | 0.247 | 0.009 |
| Direction × Scene × Experimental condition | 2.07 | 0.151 | 0.004 |
| Surround colour × Scene × Experimental condition | 0.21 | 0.892 | 0.001 |

Within each Experimental condition, Direction contrasts revealed a significant Neighbouring advantage in both the Baseline condition (0.058, *t*(466) = 2.74, *p* = 0.006) and the Suppression condition (0.145, *t*(466) = 6.82, *p* < 0.0001), with a larger effect under Suppression. Similarly, Direction effects were significant in both Indoor (0.054, *t*(466) = 2.55, *p* = 0.011) and Outdoor scenes (0.149, *t*(466) = 7.00, *p* < 0.0001), indicating a stronger Direction effect in the Outdoor scene. Overall, these results demonstrate that the influence of Direction on CCI is systematically modulated by spatial context, scene content, and experimental condition, with the strongest effects emerging when contextual cues are both chromatically salient and experimentally emphasised, see Figure 9.

**Fig. 9.**
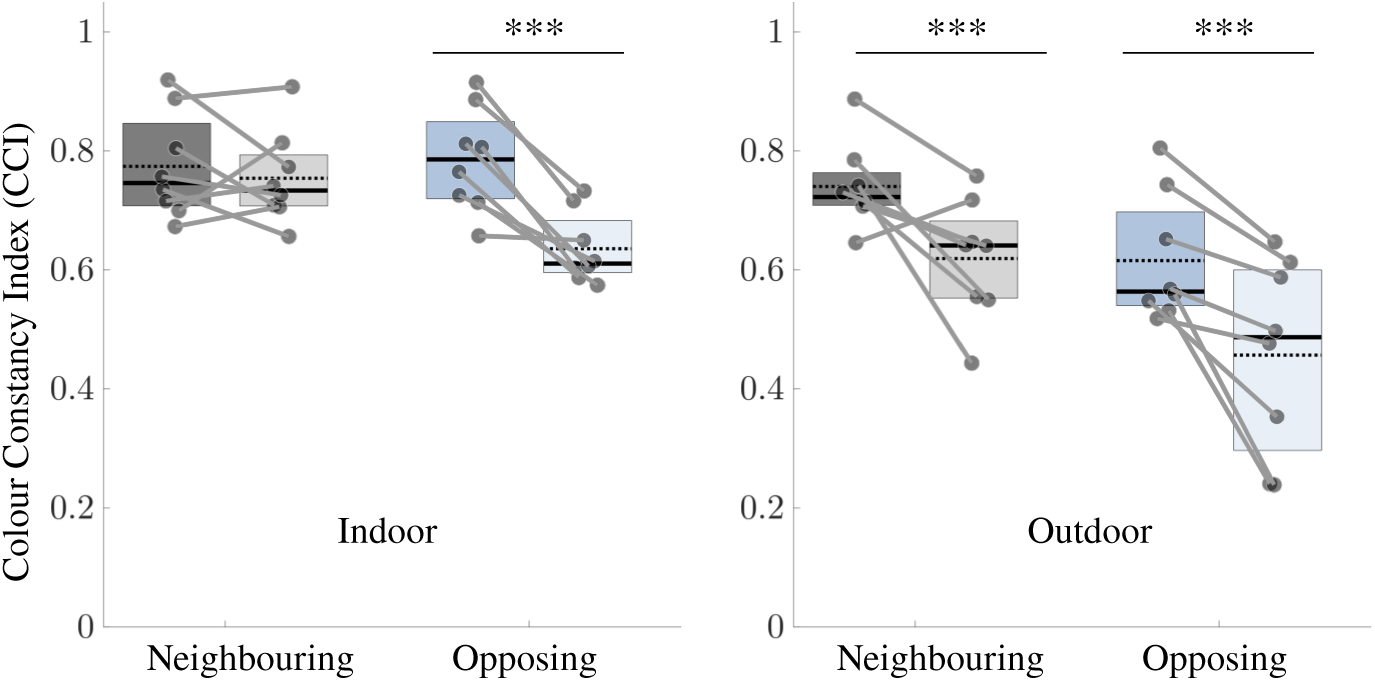
Colour Constancy Index of illuminant direction relative to the *Local Surround colour*. The left panel shows results for the *Indoor* scene, comparing *Neighbouring* and *Opposing* illuminants. For each direction, two boxplots are presented: the left box corresponds to the *Baseline* condition, and the right box to the *Surround Suppression* condition. The right panel displays the corresponding results for the *Outdoor* scene. Each dot represents a participant.

## 4. Discussion

When local surround cues were silenced, results revealed a marked decline in colour constancy in both environments, reducing it by approximately 8.5% indoors and 14.7% outdoors. Moreover, we found that constancy performance declined more sharply under illuminants opposing the local surround colour, particularly when the surround cue was silenced. Overall, these findings demonstrate that colour constancy is significantly weakened when local surround information is not available. The introduction of the concept of *Chromatic direction* helps explain the interaction observed between the illuminant colour and the local surround.

### 4.1. Consistent colour constancy with all available cues

Under the *Baseline* condition, when all cues are present, colour constancy was generally robust, especially in the *Indoor* scene. When spatial layout and local chromatic cues are consistent and reliable, the visual system can effectively discount changes in illumination. This finding is consistent with previous accounts of colour constancy that emphasise the integration of chromatic information distributed across multiple surfaces and spatial scales, rather than reliance on isolated local cues Brainard and Radonjić (2014); Foster (2011).

The *Indoor* environment, with constrained geometry and reliable relationships among surface reflectances, provides optimal conditions for estimating the illuminant and maintaining perceptual stability Radonjić and Brainard (2016). In contrast, *Outdoor* scene showed reduced constancy even in the *Baseline* condition, with significant interactions between illuminant and surround colours. Natural scenes introduce greater spatial and spectral variability, including multiple illumination components like skylight and sunlight, which make illuminant estimation more challenging Foster et al. (2006); Olkkonen et al. (2010). Our results confirm that colour constancy is shaped by contextual factors, with higher scene complexity increasing the computational demands on perceptual interpretation Foster and Reeves (2022).

### 4.2. Consequences of Silencing local surround information

Silencing local surround cues produced a marked decline in colour constancy, as evidenced by a strong main effect of *Experimental condition*, 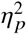 = 0.124, and a consistent decrease in CCI across scenes. In the *Indoor* scenario, silencing revealed clear illuminant-dependent biases, indicating that when local contrast information is removed, observers rely more heavily on global illumination. This shift is in line with prior evidence that local chromatic contrasts play a critical role in stabilising colour appearance Foster et al. (2006); Hou et al. (2025b); Radonjić and Brainard (2016).

In the *Outdoor* scene, silencing led to even larger performance costs and stronger interactions between illuminant and surround colours. This pattern suggests that in complex natural environments, colour constancy depends on both balanced local and global cues, both of which may be individually unreliable Brainard and Radonjić (2014). These findings align with reports that constancy deteriorates when cue eliability is reduced or when multiple illuminants are present Morimoto et al. (2024b); Olkkonen et al. (2010); Werner (2014).

### 4.3. Chromatic direction: a systematic framework for colour interactions

Introducing the factor *Chromatic direction*, distinguishing Neighbouring from Opposing illuminants relative to the surround chromaticity, revealed a powerful and systematic structure in the data. Direction exerted a strong main effect, 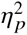 = 0.089, and interacted with Scene (Indoor and Outdoor), Surround colour (Khaki, Purple, Rose and Teal), and Experimental condition (Baseline and Surround silencing). Across analyses, Opposing illuminants consistently produced larger reductions in colour constancy performance, particularly under the *Surround silencing* condition and in the *Outdoor* scene. As illustrated in Figure 8, Δ*SE* is consistently larger under green illumination in the Outdoor scene, independent of the local surround colour. This likely reflects the fact that the scene average is already biased toward green even under neutral illumination, as shown by our previous colourimetric measurements in Gil Rodriguez et al. (2024).

This pattern supports the idea that colour constancy mechanisms are sensitive to the chromatic relationship between local context and illumination, rather than to illuminant identity alone Hurlbert and Wolf (2004). When the illuminant shifts the stimulus colour away from the chromatic axis defined by the surround, the visual system appears less able to stabilise appearance, especially when local cues are degraded. Conversely, Neighbouring illuminants tend to preserve constancy, even under the Silencing condition.

### 4.4. The influence of Scene in chromatic interactions

The *Scene* context strongly modulated the influence of *Chromatic direction* and *Surround colour*. The significant *Chromatic direction* and *Scene* interaction, 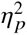 = 0.021, showed that the Outdoor scene amplifies chromatic direction effects, likely due to its greater spatial depth, reflectance diversity, and illumination complexity. In the *Indoor* scene, Direction effects were present but smaller, indicating that structured environments partially compensate for chromatic mismatches. These findings extend previous work by demonstrating that chromatic direction effects are not fixed properties of the stimulus configuration but depend on scene-level statistics. They further support the view that colour constancy emerges from adaptive cue integration tuned to natural environments Brainard and Radonjić (2014); Foster (2011).

### 4.5. Implications for computational colour constancy models

The observed correlations between chromaticity shifts and signed error Δ*SE*, together with the systematic *Direction* effects, provide important constraints for understanding colour constancy. Models that rely solely on global illuminant estimation struggle to account for the strong dependence on local surround chromaticity observed here. Instead, these results are broadly consistent with frameworks in which local adaptation, spatial comparisons, and scene interpretation jointly contribute to illuminant inference Allred and Olkkonen (2015); Foster and Nascimento (1994). Several computational models provide useful conceptual frameworks. Adaptive Surround Modulation Akbarinia and Parraga (2018) emphasises local adaptation to chromatic context, Retinex-based models Land (1986) integrate information across local and global regions, Edge-based approaches van de Weijer et al. (2007) exploit gradients and boundaries. Statistical methods such as the Shades of Grey algorithms Finlayson and Trezzi (2004) estimate the illuminant from average scene reflectance. These models illustrate how colour constancy may arise from a combination of local and global computations, where spatial structure and chromatic context modulate perception. Our findings suggest that any model aiming to capture human colour constancy should account for local surround adaptation, spatial comparisons, and scene-dependent weighting of cues to explain systematic variations with illuminant *Direction*.

## 5. Conclusions

In this study, we investigated how the interaction between illuminant and immediate local surround affects colour constancy in virtual reality environments. Using five illuminants, three along the daylight locus and two along an orthogonal chromatic direction, and four surround colours (khaki, rose, purple, and teal), we compared human performance under Baseline and Surround Silencing conditions across Indoor and Outdoor scenes.

Our findings reveal that colour constancy is highly context-dependent. In the Baseline condition, the Indoor scene exhibited robust constancy, with no significant effects of surround colour or illuminant, reflecting the visual system’s ability to integrate multiple spatial and chromatic cues under rich, all cues present environments Brainard and Radonjić (2014); Foster (2011); Radonjić and Brainard (2016). By contrast, the Outdoor scene showed a significant surround-illuminant interaction, indicating that naturalistic complexity amplifies the impact of local and global cues on colour perception Foster et al. (2006); Olkkonen et al. (2010).

The Surround silencing condition highlights the critical role of local chromatic contrast. Silencing local surrounds produced a small but reliable decrease in colour constancy, larger in Outdoor 14.7% than Indoor 8.5% environments, consistent with the greater spatial and chromatic variability of natural scenes Hou et al. (2025a,b); Morimoto et al. (2024b). In the Indoor scene, performance remained relatively stable except when the illuminant and the surround were chromatically opposed, particularly for Purple and Teal surrounds. In Outdoor scene, constancy declined broadly across illuminants, with Opposing illuminants producing the largest deficits. These patterns indicate that local surroundings provide essential contextual cues that the visual system exploits to infer illumination and stabilise colour perception Allred and Olkkonen (2015); Foster et al. (2006).

Analysis of illuminant-surround interactions revealed systematic effects: the largest signed relative errors occurred when illuminants were approximately complementary to the local surround in opponent colour space. The introduction of *Chromatic direction*, which categorises illuminants as Neighbouring or Opposing relative to the surround, clarified this effect: in both scenes, Opposing illuminants consistently produced larger constancy decrements, particularly under surround silencing, whereas Neighbouring illuminants minimally affected performance. This underscores the visual system’s sensitivity to chromatic context and its adaptive weighting of local and global cues depending on environmental complexity.

Compared to our previous work Gil Rodriguez et al. (2024), the current study found that the largest performance drop for the rose surround now occurs outdoors rather than indoors, likely due to differences in baseline design, which already incorporated a leaf matching the specific surround colour. This highlights how subtle manipulations of local context can influence constancy and suggests that cue integration strategies differ between controlled and naturalistic scenes.

These results demonstrate that colour constancy is an adaptive, context-sensitive process, strongly shaped by the interplay of illuminant, local surround, and scene complexity. Local contrast cues are particularly critical for stabilising perception under challenging illuminations, with Opposing chromatic relationships producing the most pronounced errors.

## Acknowledgments

We thank Arash Akbarinia for valuable discussions and insightful comments on the results. We also thank Dar’ya Guarnera and Giuseppe Claudio Guarnera for their contributions to the modelling and design of the 3D rendered scenes in Unreal Engine.

## Data Availability Statement

Data underlying the results presented in this paper are available in Ref.

## APPENDIX

The Appendix presents additional analyses and visualisations supporting the main results. It illustrates how the local surround and illuminant manipulations affect target appearance, and provides participants’ responses under the neutral illuminant.

### A. Illuminant and surround colours

Figure 10 illustrates the a*-b* plane of the CIELAB colour space for each of the four surround colours used in the experiment: khaki, purple, rose, and teal. Each surround colour is shown as a solid circle, indicating its chromaticity under the neutral illuminant. The chromaticities of the five illuminants (neutral, blue, yellow, green, and red) are plotted as transparent circles in each subplot. The coloured arrows indicate the direction and magnitude of the chromatic shift produced by each illuminant, with their hues corresponding to the illuminant colour.

**Fig. 10.**
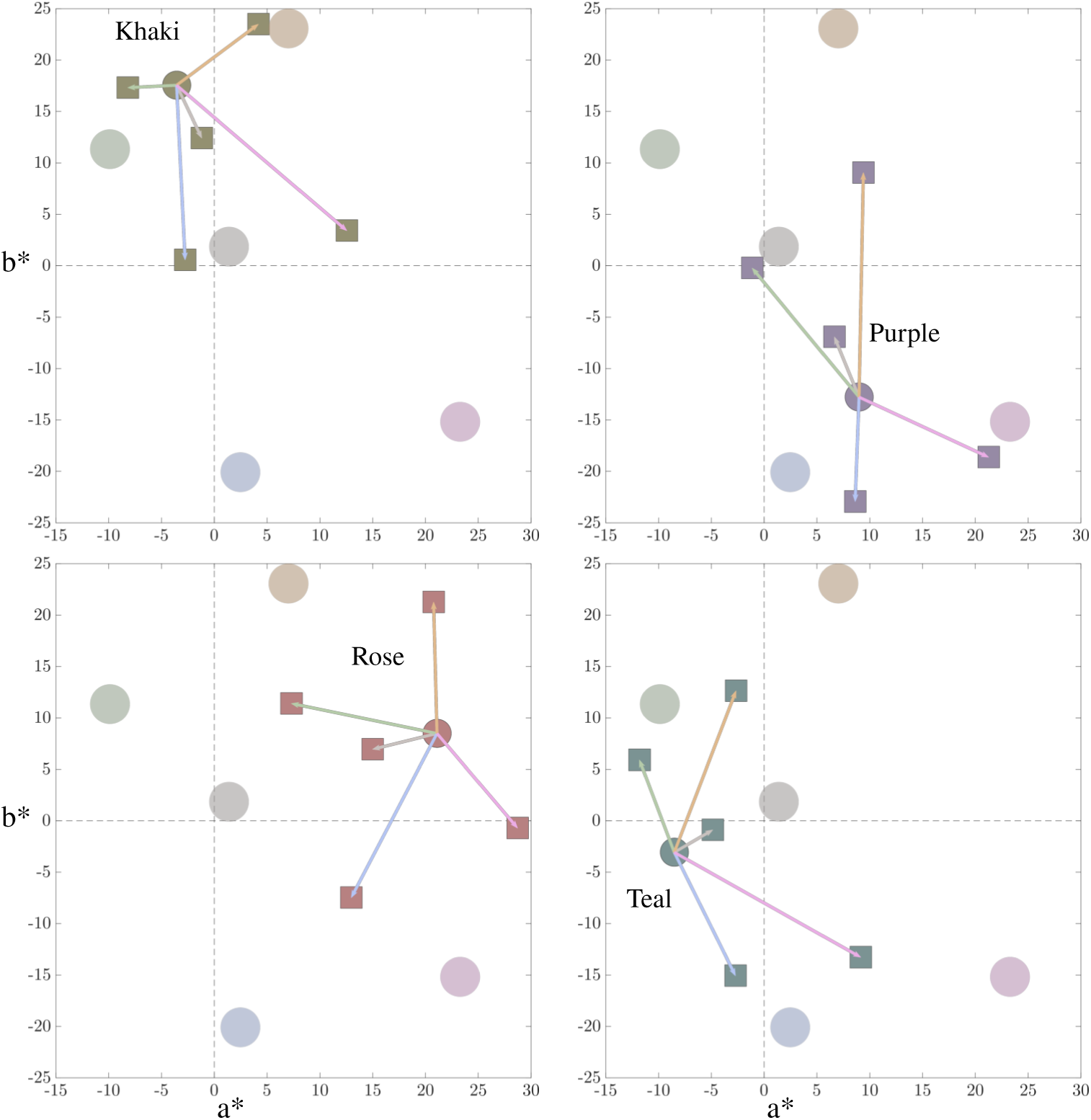
a*-b* axis of the CIELAB colour space for each Surround colour: khaki, purple, rose, and teal, shown as a solid circle. The illuminant colours are shown in each plot as transparent circles. The coloured arrows, which represent the illuminant’s colour (neutral, blue, yellow, green and red), together with the square shapes, represent the shift in colour of the Surround colour under each illuminant.

To illustrate the role of local chromatic context in our experiment, Figure 11 presents simulations of the target object and its immediate surround under all five illuminants for both the Baseline and Local Surround Silencing conditions. In the Baseline configuration, the surround changes naturally with the illumination, producing a chromatic relationship between target and context that closely resembles real-world viewing. In contrast, the Surround Silencing condition fixes the surround colour across illuminants, thereby removing this dynamic contrast cue.

**Fig. 11.**
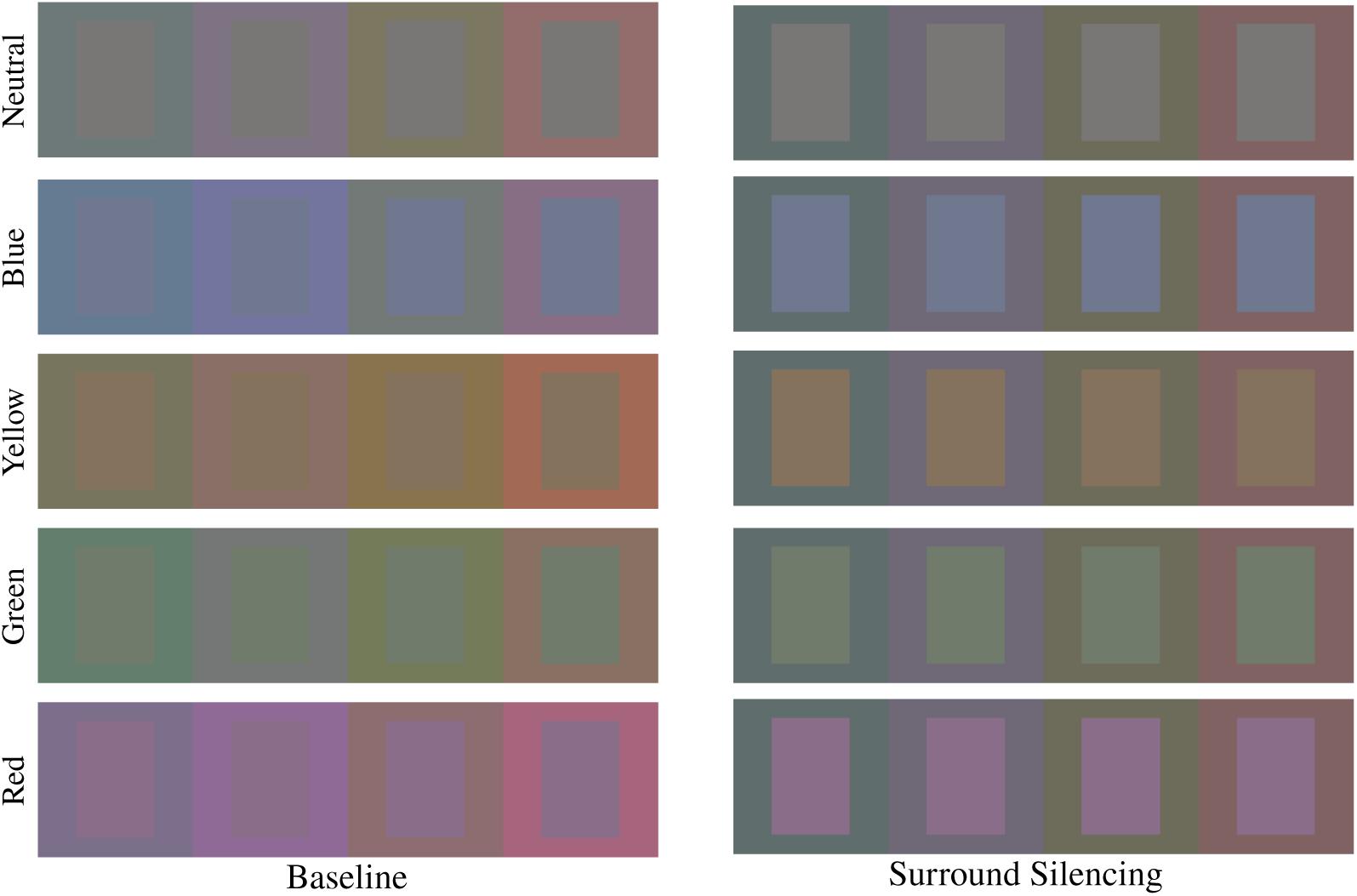
Simulated target (centre) and local surround (outer frame) for the Baseline and Local Surround Silencing conditions under five illuminants (neutral, blue, yellow, green, red). In the Baseline condition, both target and surround vary with the illuminant, whereas in the Silencing condition, the surround remains fixed while only the target changes, matching the colours of the target of the left.

### B. Results

Figures 12 and 13 present the CCI across different local surround colours: khaki, rose, purple, and teal, under all four illuminants: blue, yellow, green, and red. Each subplot contrasts the *Baseline* and *Local Surround Silencing* conditions, illustrating how colour constancy varies as a function of both scene type and surround context.

#### B.1. Pairwise Contrasts: Two-way interaction

We examined illuminant differences within each Surround colour. For the Khaki surround, none of the illuminant contrasts were significant (all |*t*| < 1.95, all adjusted *p* ≥ 0.31), with small estimates whose confidence intervals all included zero. In the Purple surround, several robust illuminant effects were observed. Blue CCIs were higher than both Green (0.200, *t*(440) = 4.74, *p* < 0.0001) and Yellow (0.194, *t*(440) = 4.69, *p* < 0.0001), Green was lower than Red (−0.132, *t*(440) = −3.12, *p* = 0.0076), and Red exceeded Yellow (0.125, *t*(440) = 3.04, *p* = 0.0076). For the Rose surround, Green was significantly lower than Red (−0.175, *t*(440) = −4.20, *p* = 0.0002), while other contrasts were non-significant or showed only weak trends. Finally, in the Teal surround, several illuminant differences were reliable: Blue CCIs exceeded Red (0.121, *t*(440) = 2.94, *p* = 0.0137) and Yellow (0.161, *t*(440) = 3.86, *p* = 0.0008), and Green was higher than both Red (0.102, *t*(440) = 2.46, *p* = 0.0425) and Yellow (0.141, *t*(440) = 3.38, *p* = 0.0039). Overall, illuminant effects were absent in the Khaki surround but pronounced in the Purple and Teal surrounds, with more selective effects in the Rose surround.

**Fig. 12.**
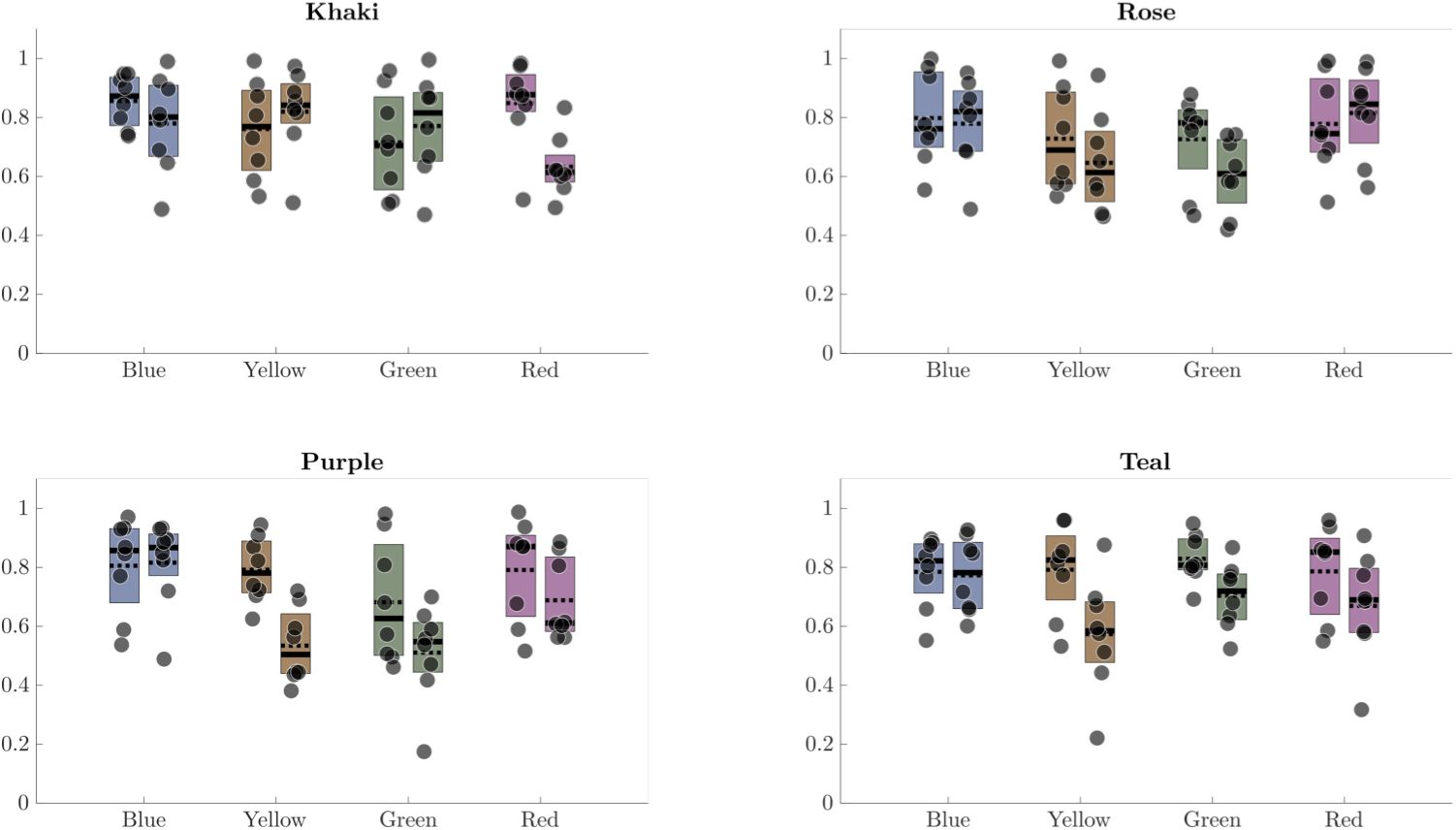
Colour constancy index for the Indoor scene for each local surround colour: khaki, rose, purple, and teal. Each local surround colour shows the Baseline along with the Surround silencing conditions under each illuminant (blue, yellow, green, red).

To assess whether illuminant effects differed between the Baseline and Surround Silencing conditions, pairwise illuminant contrasts were computed within each level of *Experimental condition* using Holm-adjusted tests. In the Baseline condition, none of the illuminant contrasts were significant (all adjusted *p* ≥ 0.95), with estimates close to zero. In contrast, several illuminant differences emerged under Surround Silencing. Blue yielded a higher CCI than Green (0.135, *t*(440) = 4.57, *p* < 0.0001) and Yellow (0.162, *t*(440) = 5.56, *p* < 0.0001), and also exceeded Red (0.091, *t*(440) = 3.13, *p* = 0.0074). Additionally, Red was higher than Yellow (0.071, *t*(440) = 2.43, *p* = 0.0467). These results indicate that illuminant-dependent differences in colour constancy are selectively enhanced when the local surround cue is silenced.

### C. Neutral Illuminant

We want to analyse how consistent participants are under the neutral illumination with each local surround colour for both Experimental conditions: Baseline and Local Surround Silencing. Under the neutral illuminant, the only difference between the Baseline and Surround Silencing conditions is that, in the Baseline, the local surround (leaf) colour interacts with light in the scene. See Section 2.4.4 for an explanation of the calculation.

Since we do not compute a CCI for the neutral illuminant, we focus on the participants’ matches in their perceptual competitors’ space, i.e., their *inferred match* Radonjić et al. (2015); Radonjić et al. (2015) (refer to Section 2.5 for more details). Since the perceptual space is one-dimensional and includes the five competitors along with participants’ responses, we rescaled all competitor positions to a fixed range from −2 to 2. In this scale, 0 corresponds to the achromatic (reference) match, while −2 and 2 represent the most bluish and most yellowish competitors, respectively. The ratios relative to the participants’ matches were maintained within this new range.

**Fig. 13.**
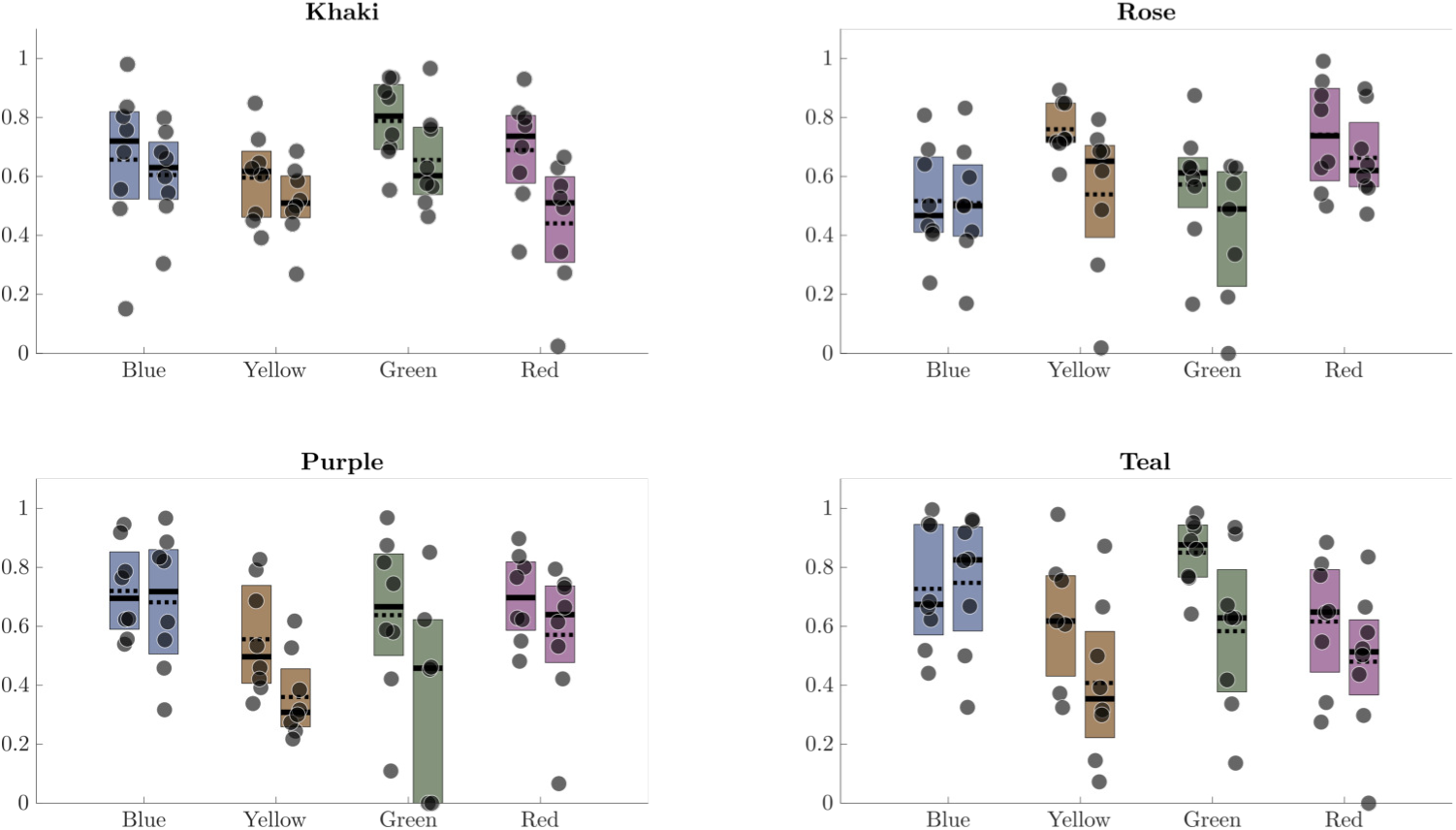
Colour constancy index for the Outdoor scene for each local surround colour: khaki, rose, purple, and teal. Each local surround colour shows the Baseline along with the Surround silencing conditions under each illuminant (blue, yellow, green, red).

We conducted a separate linear mixed-effects analysis restricted to the neutral illuminant and *inferred matches* as the responses. In this model, Surround colour, Scene, and Experimental condition were entered as fixed effects, including all two- and three-way interactions, with a random intercept for Participant.

The analysis revealed a significant main effect of Experimental condition (*F*(1, 98.6) = 4.62, *p* = 0.034), indicating that *inferred matches* differed reliably between Baseline and Silencing conditions under neutral illumination. In contrast, there were no significant main effects of Surround colour (*F*(3, 98.6) = 0.94, *p* = 0.424) or Scene (*F*(1, 39.9) = 1.18, *p* = 0.283), nor any significant two- or three-way interactions involving these factors (all *p* > 0.20).

Figures 14 and 15 (Indoor), together with Figures 16 and 17 (Outdoor), show the inferred matches under the neutral illuminant for each participant. Distinct line colours represent different surround colours. The Baseline condition is presented in Figures 14 and 16, whereas Figures 15 and 17 illustrate the Local Surround Silencing condition.

Figure 18 presents the inferred matches per participant, averaged across experimental conditions (Baseline and Surround Suppression), for both Indoor and Outdoor scenes.

**Fig. 14.**
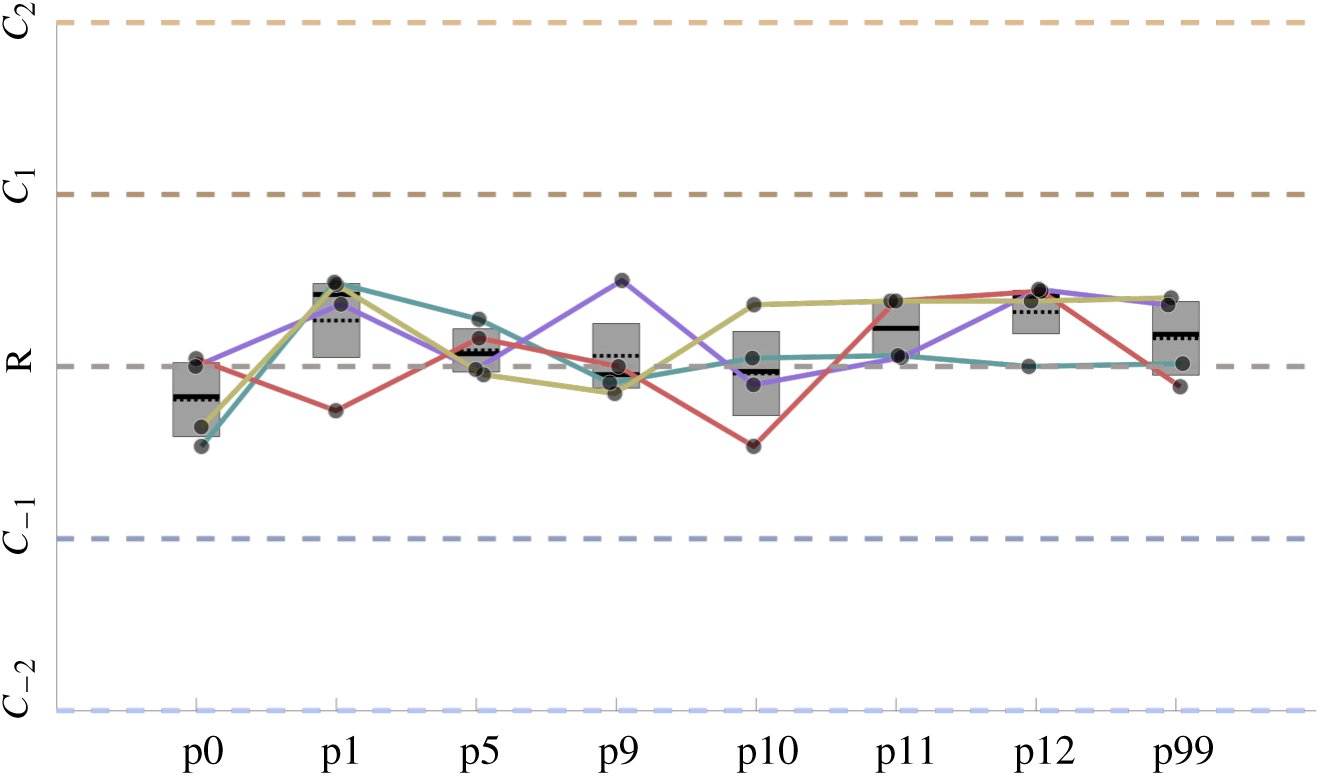
Inferred match under the neutral illuminant for each participant in the Baseline in the indoor scene. We plot together baseline vs local surround suppression for each colour surround. Rows correspond to indoor and outdoor scenes, respectively. Each boxplot represents the mean quartiles, and the error bar represents the standard deviation.

**Fig. 15.**
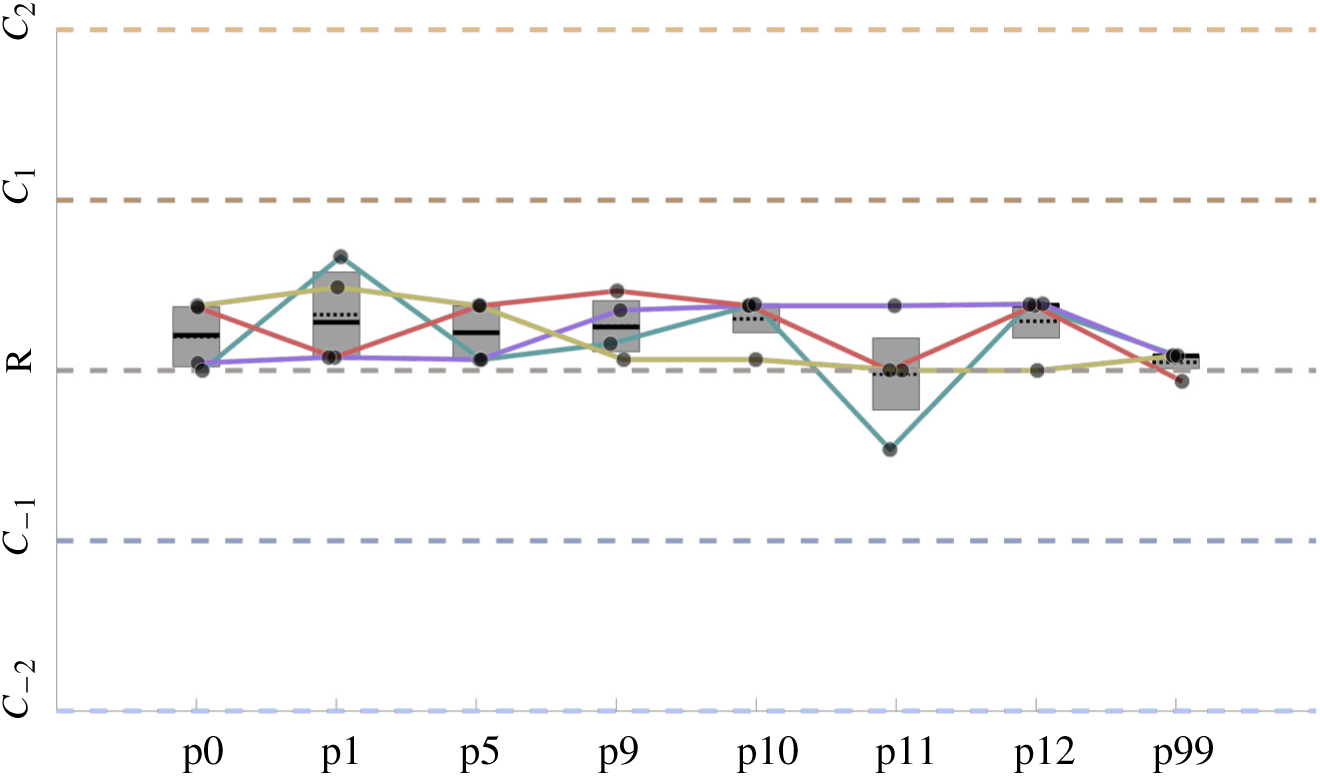
Inferred match under the neutral illuminant for each participant in the Local Surround Suppression in the indoor scene. We plot together baseline vs local surround suppression for each colour surround. Rows correspond to indoor and outdoor scenes, respectively. Each boxplot represents the mean quartiles, and the error bar represents the standard deviation.

**Fig. 16.**
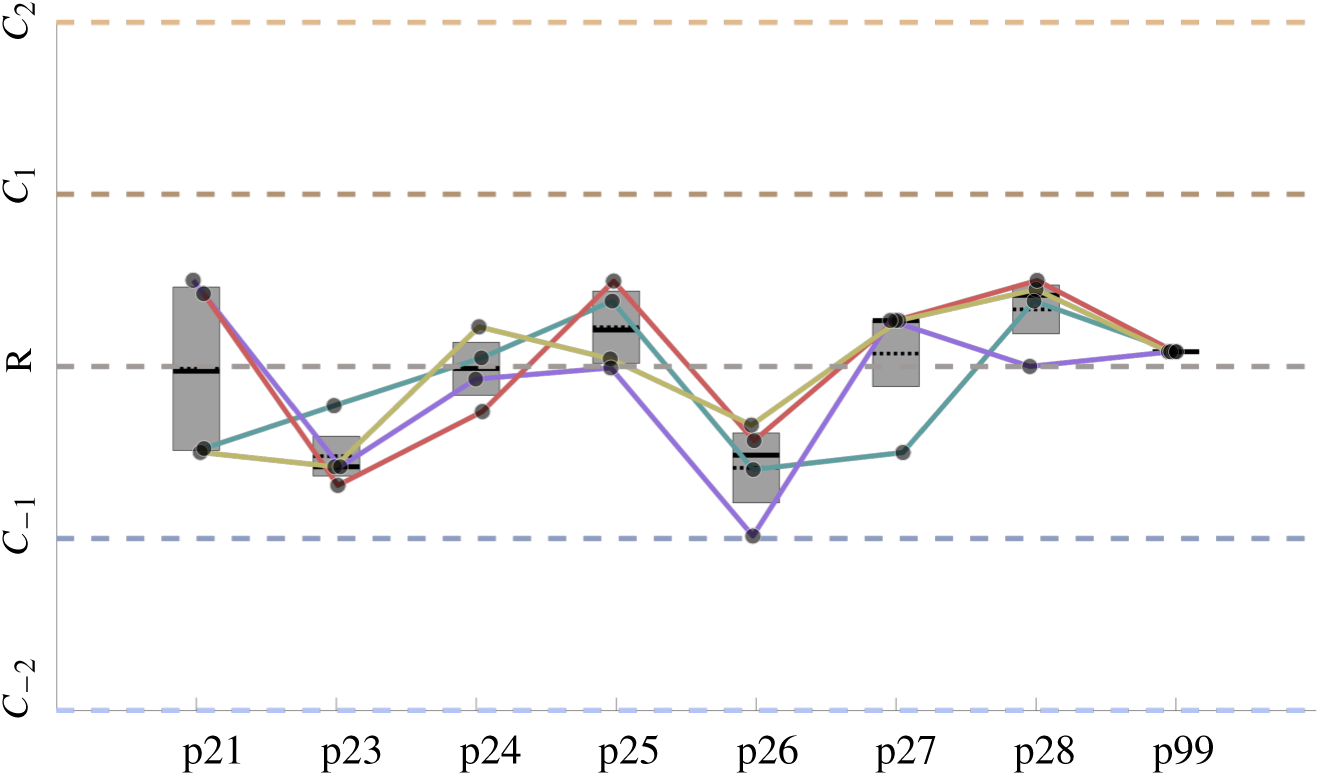
Inferred match under the neutral illuminant for each participant in the Baseline in the outdoor scene. We plot together baseline vs local surround suppression for each colour surround. Rows correspond to indoor and outdoor scenes, respectively. Each boxplot represents the mean quartiles, and the error bar represents the standard deviation.

**Fig. 17.**
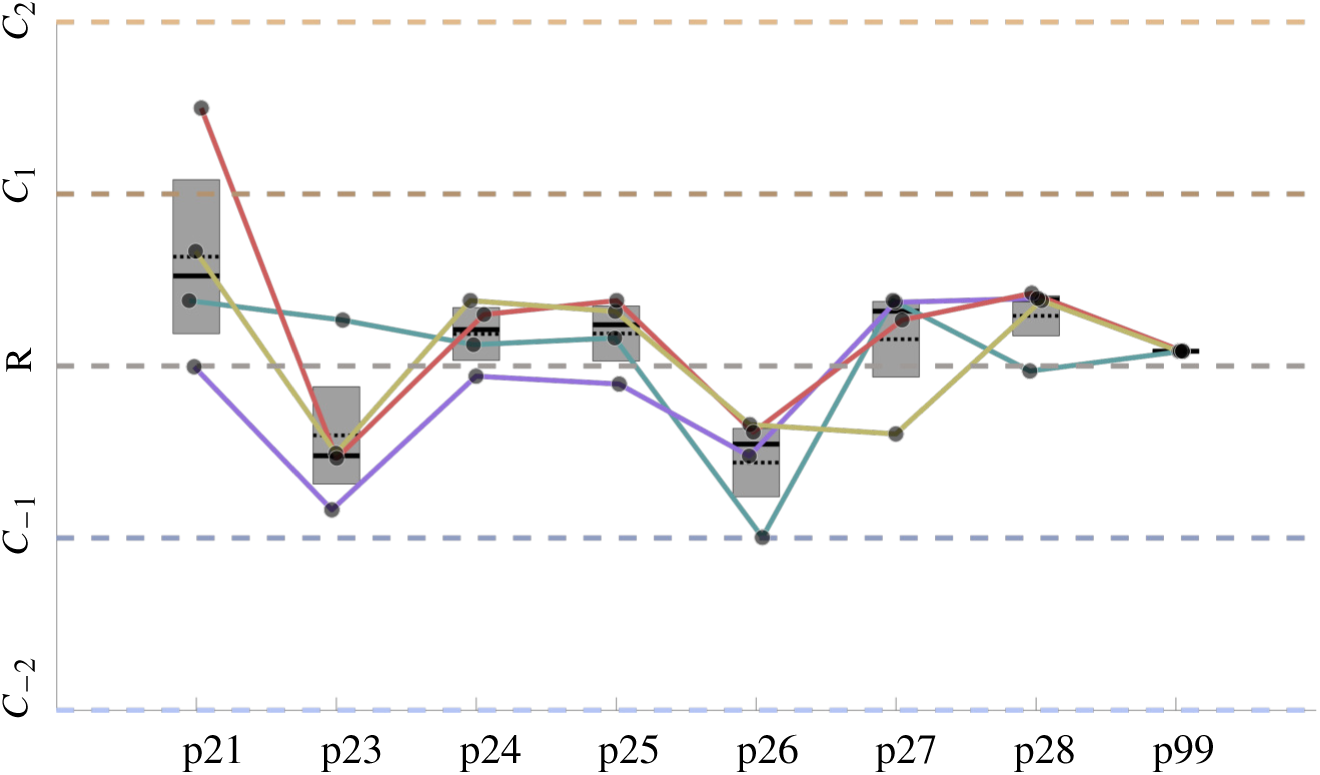
Inferred match under the neutral illuminant for each participant in the Local Surround Suppression in the outdoor scene. We plot together baseline vs local surround suppression for each colour surround. Rows correspond to indoor and outdoor scenes, respectively. Each boxplot represents the mean quartiles, and the error bar represents the standard deviation.

**Fig. 18.**
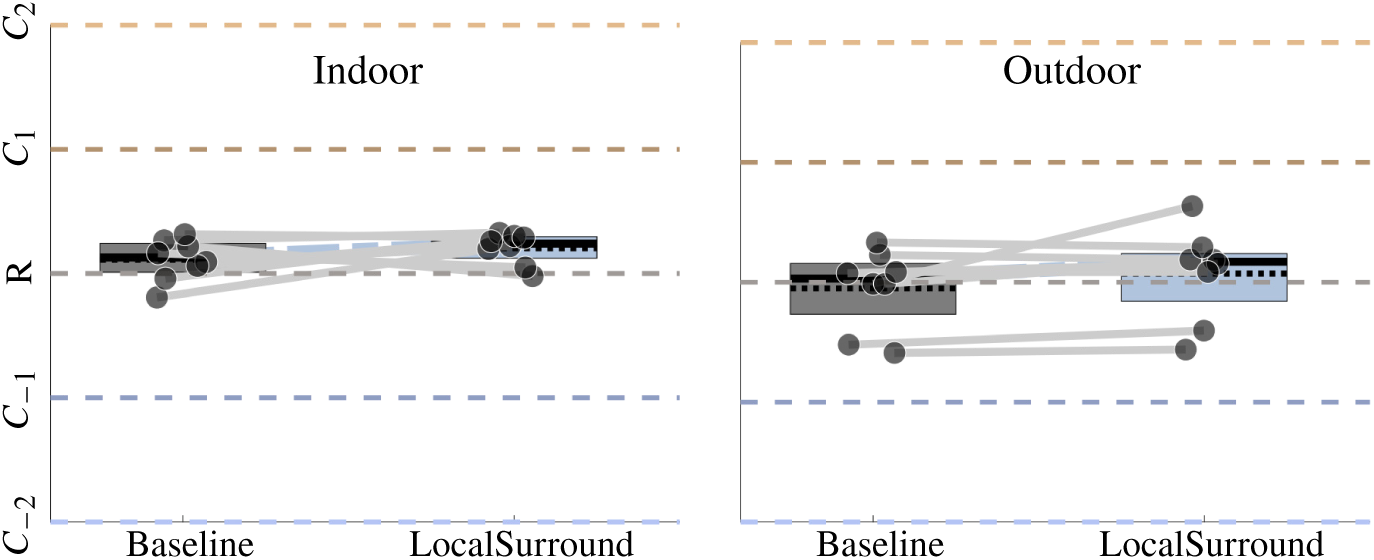
Inferred match position between baseline and local surround suppression conditions for indoor (left) and outdoor (right) scenes. It shows the average across local surround colours per participant.

